# The innate immune protein calprotectin incapacitates the bactericidal activity of β-lactam antibiotics

**DOI:** 10.1101/2025.05.14.654040

**Authors:** Amanda Z. Velez, Jana N. Radin, Emily N. Kennedy, Joshua B. Parsons, Heather M. Tong, Emma Jung, Emily Alam, Lauren C. Radlinski, Nikki J. Wagner, Vance G. Fowler, Sarah E. Rowe, Thomas Kehl-Fie, Brian P. Conlon

## Abstract

β-lactam antibiotics are widely used to treat bacterial infections, yet treatment failures frequently occur even without resistance. Here, we show that the innate immune protein calprotectin (CP), released by neutrophils and abundant at infection sites, induces tolerance to β-lactam antibiotics in *Staphylococcus aureus*. CP is a potent zinc chelator and was found to inhibit the activity of *S. aureus* autolysins, zinc-dependent enzymes essential for bacterial lysis following β-lactam-mediated inhibition of cell wall synthesis. This protection was independent of bacterial growth or metabolism and was specific to β-lactam antibiotics. Mechanistically, CP inactivated the amidase activity of Atl, the major *S. aureus* autolysin, through zinc sequestration. *In vivo*, oxacillin was significantly more effective in CP-deficient mice, demonstrating that CP reduces β-lactam efficacy during infection. These findings reveal a host-derived mechanism of antibiotic tolerance and suggest that zinc availability at infection sites may directly influence β-lactam treatment outcomes.

## INTRODUCTION

β-lactam antibiotics, such as penicillins and cephalosporins, are essential for the treatment of infections caused by a wide range of bacterial species (*1*). *Staphylococcus aureus* is the world’s leading bacterial cause of death (*2*). β-lactam antibiotics are the treatment of choice for infections caused by methicillin-susceptible *S. aureus* (MSSA) and cephalosporins such as ceftobiprole are used to treat methicillin-resistant *S. aureus* (MRSA) infection (*3, 4*). Treatment failure for *S. aureus* bloodstream infection is often attributed to antibiotic tolerance, defined as bacterial survival in the presence of an ordinarily lethal antibiotic concentration without an associated change to the minimum inhibitory concentration (MIC) (*5*). While tolerant bacteria, by definition, cannot grow in the presence of antibiotics, their increased survival necessitates prolonged antibiotic regimens and can result in treatment failure and infection recurrence once antibiotic pressure is lifted. The host conditions and factors that lead to β-lactam tolerance during infection are unknown.

β-lactams covalently bind to bacterial penicillin-binding-proteins, inhibiting the polymerization and cross-linking of peptidoglycan to reduce cell wall integrity (*6*). Autolysins are lytic enzymes functioning generally as an amidase, glycosidase, or endopeptidase, depending on the targeted bond within the cell wall (*7*). Autolysins function in opposition to penicillin-binding-proteins, maintaining the balance between cell wall degradation and synthesis. Following β-lactam inhibition of peptidoglycan biosynthesis, bacterial autolysin activity leads to cell wall degradation, lysis and death (*8-10*).

Here, we investigated the effect of a major mammalian innate immune effector, calprotectin (CP), on β-lactam treatment efficacy. CP accounts for approximately 40% of neutrophil cytoplasmic protein content and it accumulates extracellularly to mg/mL concentrations at the site of infection (*11*). A heterodimer of S100A8 and S100A9, CP tightly binds first-row transition metals, including zinc, manganese, iron, copper, and nickel (*12-15*). This activity contributes to the ability of the host to impose metal starvation on invading pathogens representing an important part of nutritional immunity (*16, 17*). Imaging studies have revealed staphylococcal abscesses are highly metal-restricted and that CP is present in high abundance (*16, 18*). In multiple models, the loss of CP increases the ability of pathogens to obtain metals and activate metal dependent processes (*17-23*). This study reveals that host-imposed zinc limitation ablates autolysin activity, dramatically reducing the killing activity of β-lactam antibiotics *in vitro* and in a murine bacteremia infection model.

## RESULTS

### The presence of calprotectin negatively impacts the efficacy of β-lactam antibiotics

As β-lactams are a first-line treatment for *S. aureus* infections and CP is present in high abundance within the abscess, how CP influences the treatment efficacy of cefazolin against the MSSA strain Newman was first examined. Following growth to mid-exponential phase in TSB-based media, a range of physiologically relevant CP concentrations was added alongside cefazolin. Interestingly, the addition of CP promoted increased staphylococcal survival in a dose dependent manner (Fig. 1A; Fig. S1A). To determine if tolerance in the presence of CP was specific to cephalosporins or generalizable across different subclasses of β-lactam antibiotics, the impact of CP on the anti-staphylococcal penicillins was next investigated. In the presence of 60 μg/mL CP, approximately 50-fold more *S. aureus* survived treatment with both oxacillin and nafcillin (Fig. 1B-C; Fig. S1B-C). To explore if changes in antibiotic efficacy were specific to the MSSA strain Newman, multiple clinical isolates of *S. aureus* (Table S1) were tested and a similar induction of antibiotic tolerance in all isolates was observed (Fig. S1D). Additionally, the presence of CP similarly increased tolerance of a clinical MRSA bacteremia isolate and the MRSA laboratory strain JE2 (Table S1) to ceftobiprole, a β-lactam approved for the treatment of MRSA bacteremia (Fig. S1E).

**Figure 1.**
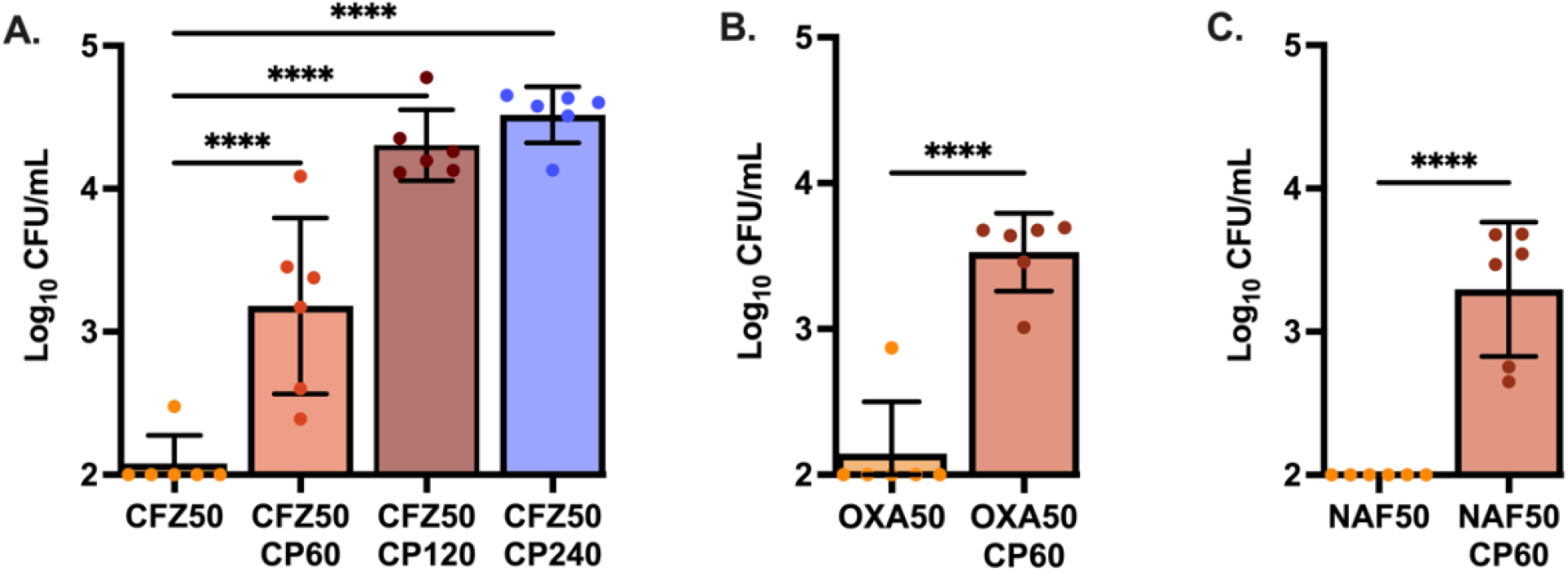
Calprotectin negatively impacts the efficacy of cell-wall acting antibiotics. *S. aureus* Newman was initially grown to mid-exponential phase in 38% TSB and 62% CP buffer. Cultures were then either treated with (A) cefazolin (CFZ50) at 50 μg/mL and CP at 60 μg/mL, 120 μg/mL, or 240 μg/mL; (B) oxacillin (OXA50) at 50 μg/mL with 60 μg/mL CP, or (C) nafcillin (NAF50) at 50 μg/mL with 60 μg/mL CP. After 24hrs treatment, an aliquot of each culture was washed and plated to enumerate surviving bacteria. Graphed are the means with error bars representing standard deviation. All experiments were completed on three separate days using two independent cultures (N=6). Statistical significance (*p*≤0.05) was determined by one-way ANOVA with Dunnett’s multiple comparison to the control group of CFZ50 (A) or by two-tailed unpaired t-test (B-C).

To examine the possibility that general protein binding was driving antibiotic inefficacy, antibiotic killing in bovine serum albumin (BSA) at similar protein concentrations to CP was examined.

Our results showed that BSA had no impact on survival post oxacillin treatment (Fig. S1F). Additionally, the observed increased survival against antibiotics in the presence of CP was not associated with a change in the minimum inhibitory concentration (MIC) of the antibiotic (Table S2), demonstrating that antibiotics remain unbound and capable of inhibiting their target in the presence of CP. Together, our results show that CP dramatically increases survival against the main class of antibiotics used in the treatment of *S. aureus* infections, with no detectable change to the MIC. Furthermore, this effect was observed across multiple β-lactam antibiotics and across multiple *S. aureus* strains, both MSSA and MRSA.

### Zinc limitation induces antibiotic tolerance in a mechanism independent of target site activity or metabolic state

Because CP sequesters zinc and manganese via two distinct metal-binding sites located at the dimer interface of the S100A8 and S100A9 subunits: S1, a His_6_ motif and S2, a His_3_Asp motif. A CP variant that lacks both metal binding sites and cannot bind transition metals, was examined for its capacity to induce antibiotic tolerance (*12*). The mutant protein was unable to induce tolerance to oxacillin, strongly implicating metal limitation as the driver of β-lactam tolerance. (Fig 2A; Fig. S2A).

**Figure 2.**
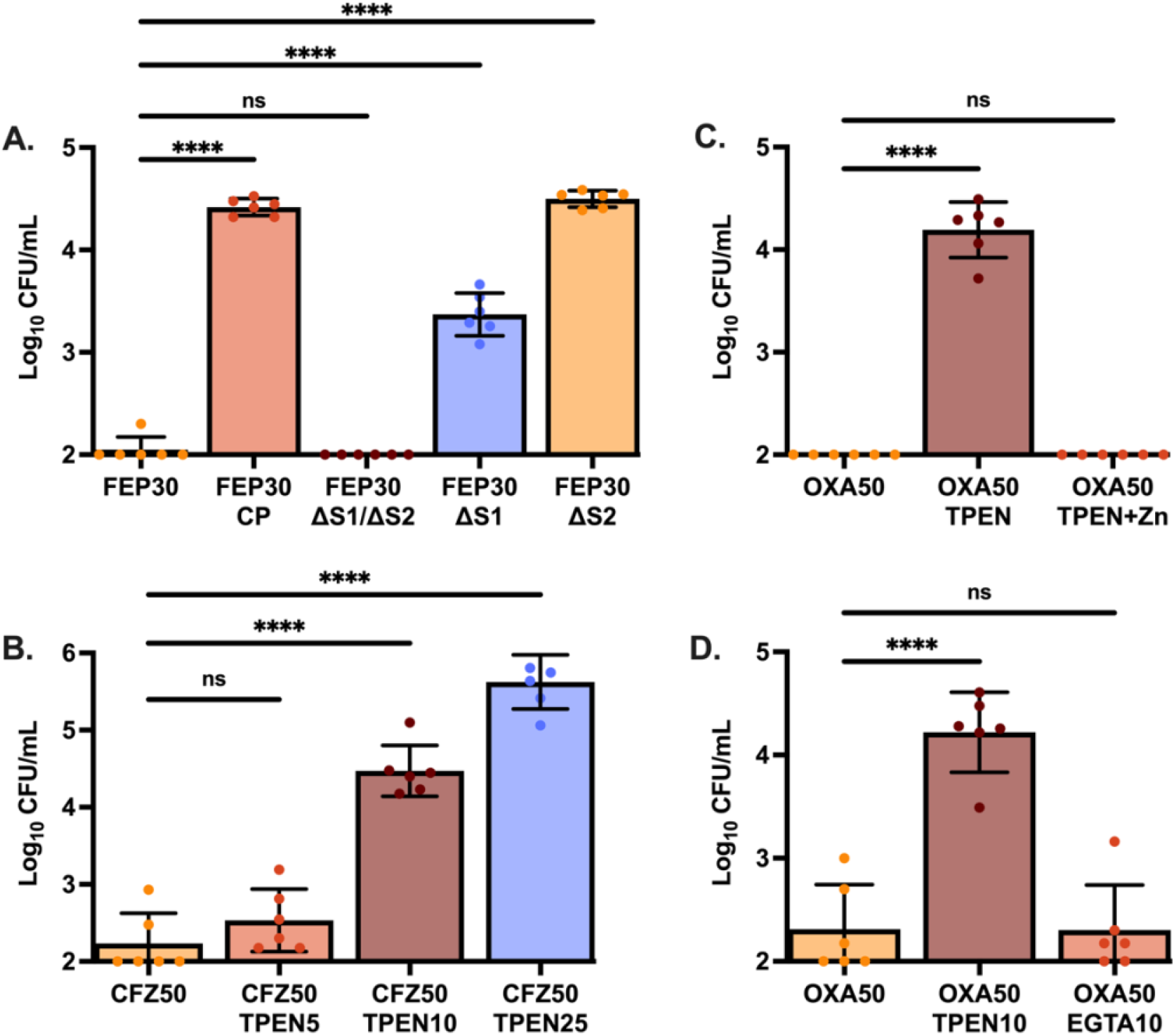
Zinc limitation induces antibiotic tolerance. *S. aureus* Newman was grown to mid-exponential phase in 38% TSB and 62% CP buffer. Cultures were then treated with (A) cefepime at 30 μg/mL (FEP30) and either wildtype CP (240 μg/mL), ΔS1/ΔS2 (240 μg/mL), ΔS1 (480 μg/mL), or ΔS2 (480 μg/mL). Single site mutant concentrations were doubled as they have half the binding capacity of wildtype CP; (B) cefazolin at 50 μg/mL (CFZ50) and increasing concentrations of TPEN, 5 μM to 25 μM; (C) oxacillin at 50 μg/mL (OXA50) and 10 μM TPEN with and without the addition of 50 μM ZnSO_4_; or (D) oxacillin at 50 μg/mL (OXA50) and 10 μM TPEN or 10 μM EGTA. Data graphed represents the mean survival 24 hours post treatment with error bars showing standard deviation. Each experiment was completed on three separate days using two independent cultures (N=6). Statistical significance (*p*≤0.05) was determined by one-way ANOVA with Dunnett’s multiple comparison to survival in the absence of chelator.

To identify which metal was responsible for reducing antibiotic efficacy, single site disruptions for metal binding (ΔS1 and ΔS2) were examined (*12*). S1 binds both zinc and manganese while S2 binds only zinc. If zinc sequestration is driving β-lactam tolerance, then the single mutants will retain the ability to induce tolerance, whereas if manganese limitation is driving the phenotype, the S1 mutation alone should be sufficient to ablate the induction of tolerance. Comparison between the two site mutants suggested CP decreases antibiotic efficacy primarily through the limitation of zinc as both variants significantly increase bacterial survival post cefepime treatment (Fig. 2A; Fig. S2A).

To confirm that zinc limitation induces β-lactam tolerance, antibiotic killing in the presence of the potent zinc chelator, N,N,N’,N’-tetrakis(2-pyridinylmethyl)-1,2-ethanediamine (TPEN) was examined. Similar to CP, the addition of TPEN resulted in decreased efficacy of β-lactam antibiotics in a dose-dependent response, culminating in a 1000-fold increase in *S. aureus* survivors relative to the metal replete control (Fig. 2B; Fig. S2B). Furthermore, supplementation of the media containing both TPEN and exogenous ZnSO_4_ was sufficient to restore antibiotic efficacy (Fig. 2C; Fig. S2C). When EGTA, a potent chelator of calcium with limited affinity for zinc, was used, no change to antibiotic killing kinetics as compared to the chelator absent control was observed (Fig. 2D; Fig. S2D). Together, these data demonstrate that zinc limitation leads to the development of tolerance to β-lactam antibiotics.

Antibiotic tolerance has frequently been associated with slower growth, decreased metabolism and reduced target activity. However, the effect of CP on β-lactam survival was observed at a concentration two-fold lower than the IC50 for CP in these conditions (*12*). Consistent with this, the CFU recovered in the presence of 60 μg/mL CP was similar to media lacking CP (Fig. 3A). This suggests that CP-induced tolerance is not a direct result of slowed growth. To examine cell-wall biosynthesis activity, the fluorescent d-amino acid (HADA) which accumulates at sites of cell wall synthesis and a cell surface carbohydrate stain (wheat germ agglutinin conjugated to Alexa Fluor 488), were used to visualize regions of active cell wall synthesis. These results showed that the addition of CP did not result in the inhibition of cell wall synthesis, as seen by the accumulation of HADA signal comparable to that of untreated cells (Fig. 3B). This suggests that zinc limitation induces antibiotic tolerance in a mechanism independent of target site activity.

**Figure 3.**
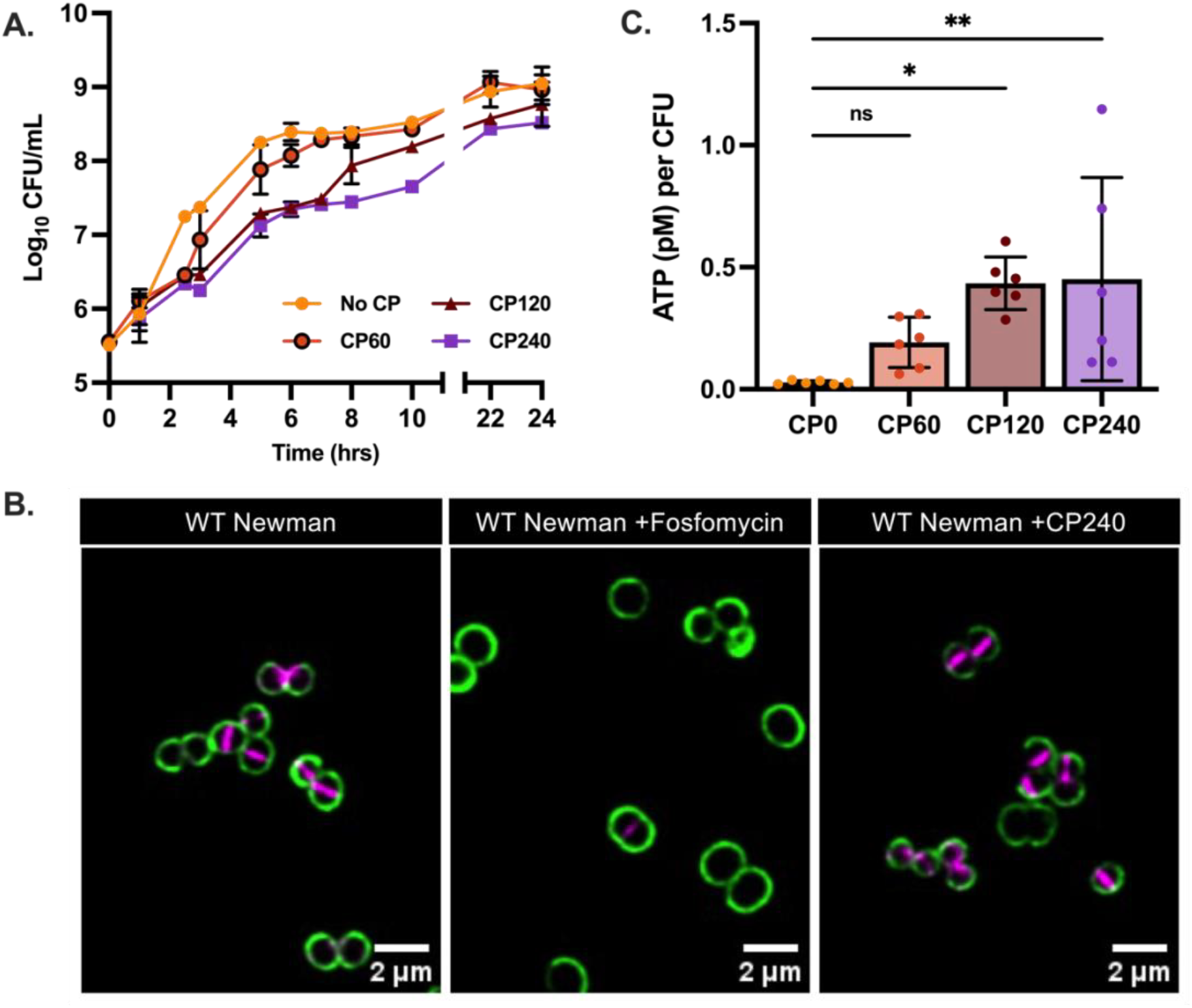
Zinc limitation induces antibiotic tolerance in a mechanism independent of target site activity or metabolic state. (A) Growth curve of *S. aureus* Newman in TSB containing CP at concentrations ranging from 0-240 μg/mL. This experiment was completed on two separate days using two independent cultures (N=4). (B) *S. aureus* Newman was grown in 38% TSB and 62% CP buffer to mid-exponential phase and then either left untreated or treated with CP (240 μg/mL) or fosfomycin (50 μg/mL) for 1 hour. Cells were then stained with HADA followed by WGA-488 and representative cells of the population are presented for each condition. (C) At ∼10^6^ CFU/mL, ATP levels of cells growing in the presence of CP (0-240 μg/mL). This experiment was completed on three separate days using two independent cultures (N=6). Statistical significance (*p*≤0.05) was determined by one-way ANOVA with Dunnett’s multiple comparison to the survival in the absence of CP (CP0).

To examine the energy state of the cell in the presence of CP, the BacTiter-Glo Microbial Cell Viability Assay was utilized, measuring ATP levels through a luminescent reaction with the luciferase enzyme. Measurement of cellular ATP levels with and without the addition of CP showed no reductions in the amount of ATP calculated per CFU (Fig. 3C), instead ATP levels increased, suggesting that while CP appears to impact the metabolic state of the cell, CP induced tolerance is independent of a reduction in metabolic activity.

### Bacterial autolytic activity is decreased by the zinc-limiting activity of calprotectin

Because growth, target site activity or metabolic state are not decreased in the presence of CP, tolerance must be induced through an alternative mechanism. β-lactam antibiotics inhibit cell wall synthesis by covalently binding to transpeptidases or penicillin-binding-proteins. This inhibition prevents the cross-linking of peptidoglycan chains, weakening the cell wall. The resulting cell lysis is thought to be mediated by autolysins, enzymes that degrade the cell wall. Supporting this model, multiple studies have shown that deletion of autolysin genes enhances bacterial survival following β-lactam treatment (*8-10*). In line with previous research, deletion of autolysins, Atl and LytM, leads to ablation of the bactericidal activity of oxacillin (Fig. S2E). The genome of *S. aureus* encodes for multiple autolysins, four of which require zinc as a cofactor for catalytic activity, including Atl, the primary *S. aureus* autolysin. To examine the requirement of zinc for autolytic activity, parallel zymography was performed in the presence and absence of TPEN. Comparison of gels with and without TPEN, showed that the addition of TPEN inhibited Atl-mediated cell wall clearance (Fig. S3).

To examine if CP can reduce autolysis through zinc limitation, the supernatant of bacterial cultures was combined with heat-killed *S. aureus* cells and autolytic activity was assessed. Active autolysins will lyse heat-killed *S. aureus*, resulting in a decrease in OD_600_ over time. In the presence of CP, complete loss of autolytic activity was observed as represented by little change in OD (Fig. 4A). When CP variants were used, the ΔS1/ΔS2 mutant, but not the ΔS1 or ΔS2 mutant, both which retain zinc binding, resulted in complete recovery of autolytic activity (Fig. 4B). Since both sites can bind zinc, recovery of autolytic activity for only ΔS1/ΔS2 implies that zinc limitation is responsible for the lack of autolytic activity in the presence of CP.

**Figure 4.**
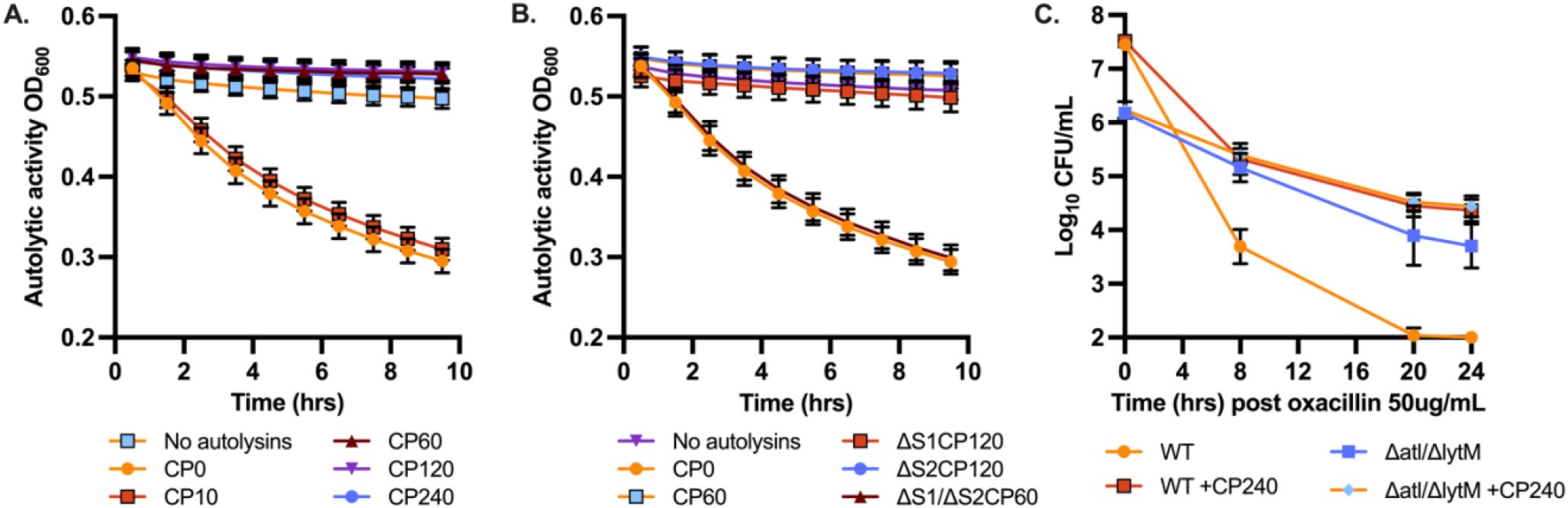
Autolytic activity is decreased by the zinc limiting activity of calprotectin. (A) *S. aureus* Newman was grown in 38% TSB and 62% CP buffer overnight prior to pelleting the cells and combining the supernatant with heat-killed *S. aureus* RN4220 cells and increasing concentrations of CP (0-240 μg/mL). (B) Cell supernatant was combined with heat-killed *S. aureus* RN4220 cells in the presence of wildtype CP (60 μg/mL), ΔS1/ΔS2 (60 μg/mL), ΔS1 (120 μg/mL), or ΔS2 (120 μg/mL). (C) *S. aureus* Newman wild-type or Δ*atl*/Δ*lytM* was grown for 3 hours and then challenged with oxacillin at 50 μg/mL with and without 240 μg/mL CP. Surviving cells were enumerated at 8, 20, and 24 hours.

**Figure 5.**
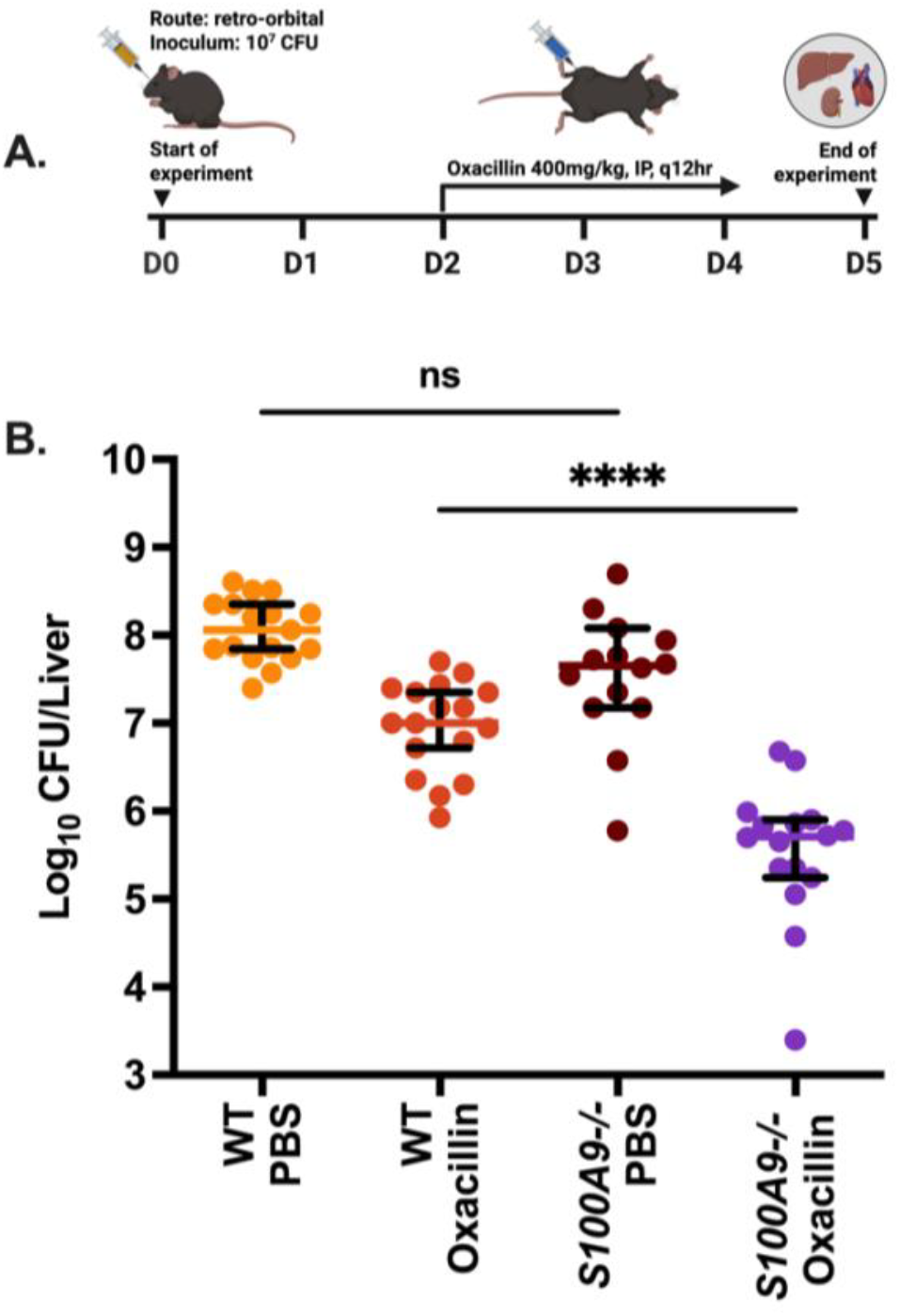
The presence of calprotectin negatively impacts the efficacy of cell-wall acting antibiotics *in vivo*. (A) Timeline of infection for the murine bacteremia model. Wild-type and S100A9^-/-^ C57BL/6 mice were infected retro-orbitally with 10^7^ CFU of *S. aureus* Newman. After 48hrs, oxacillin intraperitoneal treatment at 400 mg/kg administered every 12 hours was initiated and continued over three days. At five days post-infection, mice were euthanized and select internal organs were homogenized, diluted, and plated to determine bacterial burden. (B) Surviving bacteria within the liver of infected mice at day five post infection. Data graphed is the median with 95% CI. This experiment was completed twice with each data point representing an individual mouse. Statistical significance was determined by one-way ANOVA with Tukey’s multiple comparisons test. Results are considered significant if *p*≤0.05.

To determine if autolytic activity was responsible for the induction of tolerance in the presence of CP, mutations were introduced into the genes encoding the zinc-dependent autolysins, *atl* and *lytM* (Fig. 4C). Loss of two zinc-dependent autolysins (Δ*atl*/Δ*lytM)* clearly reduced the differences in killing observed with the addition of CP. This suggests that the majority of the tolerance observed in the presence of CP is due to inhibition of autolysin activity. To visualize potential changes in the cell wall of *S. aureus* because of decreased autolytic activity in the presence of CP, cell wall structure was also examined by transmission electron microscopy.

The addition of CP showed noticeable changes to the structure of the cell walls after 2 hours (Fig. S4). Comparison between cross-sections showed regions of increased peptidoglycan for CP-treated cells (black arrows). This is reminiscent of previous observations for Δ*atl S. aureus*, where absence of Atl activity leads to thread-like peptidoglycan interconnections between cells (*24*).

### The presence of calprotectin negatively impacts the efficacy of cell-wall acting antibiotics during bacteremia

As CP is a heterodimeric S100 protein formed by association of monomers S100A8 and S100A9, mice that are *S100A9-/-* are essentially deficient in CP (*25*). To determine the impact of CP on the *in vivo* activity of cell-wall targeting antibiotics, both wildtype and *S100A9-/-* mice were infected with *S. aureus* Newman retro-orbitally and then treated with oxacillin (Fig. 6A). Oxacillin showed significantly improved bacterial clearance in the liver of CP-deficient mice with approximately 20-fold less survivors relative to wild-type mice (Fig. 6B).

These effects were not observed in the kidney and heart tissue where no significant differences in oxacillin treatment between wild-type and *S100A9-/-* mice were observed (Fig. S5A-B).

Together these results demonstrate that CP induces potent tolerance to β-lactam antibiotics in *S. aureus* through inactivation of zinc-dependent autolysins, resulting in reduced β-lactam efficacy *in vitro* and *in vivo*.

## DISCUSSION

The treatment of *S. aureus* infections is characterized by a high incidence of failure, even in cases involving drug-susceptible strains (*26, 27*). Several studies have shown that antibiotic efficacy is dramatically modulated by the host (*28-32*). Understanding how antibiotics function within the infectious environment is key to preserving and enhancing their efficacy. This study investigated how CP, one of the major proteins of the innate immune system, influences the efficacy of β-lactam antibiotics against *S. aureus* infection. Our results show that CP-driven zinc chelation induces tolerance to β-lactam antibiotics, through the inhibition of zinc-dependent autolysins. Furthermore, CP significantly inhibits oxacillin efficacy in a mouse model of bacteremia.

CP is an abundant protein within the neutrophil cytosol and at sites of infection the influx of neutrophils leads to the accumulation of CP in excess of 1 mg/mL (*11*). The metal-binding ability of CP plays a key role in nutritional immunity, the process by which essential nutrients for microbial survival are limited by the host. While CP binds multiple first row transition metals tightly (*12-15*), the current results strongly support that zinc limitation induces tolerance to β-lactam antibiotics. The contribution of CP to metal limitation varies by tissue. However, CP is not the only zinc binding protein found at sites of infection, with humans producing both S100A7, secreted by keratinocytes, and S100A12, secreted by neutrophils, as well as other unidentified mechanisms for restricting zinc availability (*16, 18, 33, 34*). These zinc withholding mechanisms could all contribute to reduced β-lactam efficacy within the host, affecting the impact that loss of CP has on antibiotic tolerance. Despite these additional mechanisms, loss of CP does increase the availability of zinc (*22, 23*), particularly in the liver, as evidenced by the increased virulence of an *A. baumannii* zinc-transporter mutant in the liver of CP-deficient mice (*17*). This suggests that the increased efficacy of β-lactams observed in the liver of CP-deficient mice in this study is driven by increased zinc availability.

Even in the presence of high levels of CP, *S. aureus* is well-equipped to transport zinc and maintain sufficient levels for intracellular functionality (*25, 35*). However, extracellular zinc-dependent enzymes are vulnerable to CP sequestration of zinc. In support of this, CP can potently inhibit the zinc-dependent protease activity of *P. aeruginosa* extracellular virulence factors LasA and LasB (*36*). Additional *S. aureus* metalloproteases may also be vulnerable to metal limitation by CP. The extracellular zinc metalloprotease, aureolysin, which cleaves host proteins and antimicrobial peptides, functions as an important virulence factor for *S. aureus* (*37*), suggesting that CP may influence *S. aureus* pathogenesis as well as antibiotic susceptibility.

Numerous autolysins over a broad range of Gram-positive and Gram-negative species are extracellular metalloenzymes requiring zinc for autolytic activity, including *Clostridioides difficile* Cwp6, *Streptococcus pneumoniae* LytA, *Helicobacter pylori* Csd2, and *Escherichia coli* AmiA/B/C, among many others (*38-41*). Thus, the role of zinc limitation in altering cell wall homeostasis remains to be investigated in depth and may prove to be a major determinant of antibiotic susceptibility for multiple pathogens.

Additional zinc-dependent extracellular enzymes capable of influencing antibiotic treatment outcomes include metallo-β-lactamases. Previous research has shown that the addition of TPEN was sufficient to restore antibiotic susceptibility below the clinical breakpoint for a carbapenem resistant clinical isolate of *A. baumannii* expressing a zinc-dependent carbapenemase (*17*). Inactivation of extracellular zinc-dependent enzymes therefore likely represents an important function of CP during infection, albeit with major negative consequences for β-lactam efficacy.

The levels of transition metals available to pathogens at the site of infection is also dependent on dietary intake and can be manipulated by diet modification (*42*). Increased manganese in the diet has been found to increase the availability of this metal during *S. aureus* infection (*43*). Therefore, it is interesting to consider the possibility that a high zinc diet could increase autolytic activity within the abscess, potentially improving β-lactam bactericidal activity at these difficult to treat infection sites.

In summary, these results show that CP dramatically increases survival against the main class of antibiotics used in the treatment of *S. aureus* infections. This study represents the first description of a host-induced state of antibiotic tolerance through the specific inactivation of a bacterial enzyme. While tolerance has broadly been understood in the context of metabolic inactivation or growth inhibition, our research suggests that a myriad of other host-pathogen interactions may have major impacts on antibiotic activities. These interactions remain underexplored and may represent a crucial determinant of antibiotic susceptibility that could be targeted to improve antibiotic treatment outcomes in patients.

## METHODS

### Growth conditions, strains, and reagents

For all experiments unless otherwise clarified, *S. aureus* was grown overnight in 5mL of Tryptic Soy Broth (TSB) and then diluted the following day in 3mL of 38% TSB and 62% of a high-calcium buffer (20mM Tris-HCl, 3mM CaCl_2_, 100mM NaCl, pH 7.5) (CP buffer). All bacterial growth was completed using a roller drum at 37 °C in closed cap 15mL conical tubes. Construction of Δ*atl/ΔlytM S. aureus* was completed using single mutants from the Nebraska transposon Mutant Library (NTML) and the NTML Genetic Toolbox allelic exchange system for the replacement of resistance markers, as previously described (*44*). Mutations were confirmed by antibiotic resistance to kanamycin/erythromycin and by PCR, using primers CCCTGCTATTGTCCAACCAA for *atl* and AATTAACAGCAGCAGCGATTG for *lytM*, as well as the transposon specific primers as provided by NTML. Recombinant CP lacking cysteine residues to prevent disulfide bond formation was expressed and purified as previously described (*45*). To avoid use of an ampicillin resistance cassette which encodes for a β-lactamase that was found to purify alongside CP, the expression vector was modified to include only a kanamycin resistance cassette.

### Timed kill curve assay

Overnight cultures of *S. aureus* were diluted 1:1000 and incubated to reach mid-exponential phase, between 2-3 hours depending on the strain of *S. aureus* utilized in each experiment. Following determination of the starting bacterial burden (0hr timepoint), cultures were then treated simultaneously with antibiotic and either CP, BSA, TPEN, or EGTA, at concentrations specified in each figure legend. Calcium was directly added to purified aliquots of CP at 3mM CaCl_2_ prior to use. All cultures were then monitored for survival over a period of 24hrs and the CFU/mL at timepoints 0hr, 8hr, 20hr, 24hr were examined. At each timepoint, a 100μL aliquot was washed twice in PBS, diluted, and plated on Tryptic Soy Agar (TSA) plates. Colony-forming units (CFU) were enumerated following incubation at 37 °C overnight and graphed as bacterial survival over time.

### Cell wall imaging using fluorescence microscopy

Imaging of the bacterial cell wall using fluorescence microscopy was accomplished as previously described (*46*). *S. aureus* Newman was back diluted 1:1000 and incubated to reach 10^7^ CFU/mL. Select cultures were then left untreated or treated with CP at 240 μg/mL or fosfomycin at 50 μg/mL for 1 hour. Cell walls were then stained with 250μM fluorescent D-amino acid HADA (Tocris Bioscience) for 5 minutes, followed by washing and staining with wheat germ agglutinin conjugated to Alexa Fluor 488 (WGA-488, Invitrogen) at 2μg/mL for an additional 5 minutes. Cells were then washed and fixed with 4% paraformaldehyde and 2μL pipetted between a glass cover slip and an agarose pad. Z-stack images of representative cells were then acquired using the Zeiss LSM900 microscope using the 63x/1.4 oil plan apo objective and the Zeiss 705 camera. Images were deconvolved using AutoQuant and processed with identical display adjustments using FIJI.

### Growth curve assay

Overnight cultures of *S. aureus* Newman were diluted 1:1000 in the presence of increasing concentrations of CP ranging from 0-240μg/mL. Over a period of 24hrs, 10μL of culture was diluted and plated on TSA to determine CFU/mL.

### Measurement of bacterial ATP

The BacTiter-Glo Microbial Cell Viability Assay (Promega) was utilized to measure ATP levels according to the manufacturer’s instructions. Briefly, overnight cultures of *S. aureus* Newman were diluted 1:1000 in the presence of CP (0-240μg/mL) and at around 10^6^ CFU/mL, an aliquot of culture was combined 1:1 with BacTiter-Glo Reagent in a white opaque-walled 96-well plate. For each experiment, in order to calculate ATP per cell, an ATP standard curve was also generated with concentrations ranging from 1000nM to 0.01nM, and the CFU of the culture was determined by dilution and plating. Results were adjusted to account for background signal and calculated ATP concentrations were then divided by the total number of cells per well.

### Autolysin activity assay

Overnight cultures of *S. aureus* Newman grown in 38% TSB and 62% CP buffer were pelleted and the supernatant containing autolysins was collected. Aliquots of supernatant were then combined with CP at a range of concentrations (0-240μg/mL) or combined with metal binding site disruptions, ΔS1/ΔS2 at 60μg/mL, ΔS1 at 120μg/mL, or ΔS2 at 120μg/mL. For comparison, single site mutant concentrations were doubled as they have half the binding capacity of wildtype CP. Following a 1hr incubation, 30μL of supernatant was then combined with 70μL of heat-killed *S. aureus* RN4220 resuspended in CP buffer in a 96-well plate. Over the course of 10 hours, the OD_600_ was measured and plotted against time to examine autolytic activity.

### Murine bacteremia model

*In vivo* infections were completed at the University of Illinois Urbana-Champaign following approval by the Institutional Animal Care and Use Committee (IACUC). Female wild-type C57BL/6 mice and CP-deficient mice (S100A9^-/-^ C57BL/6) were infected retro-orbitally with 10^7^ CFU of *S. aureus* Newman. The infection inoculum was prepared by back-diluting overnight cultures 1:100 in 5mL of TSB. Cultures were then incubated using a roller drum at 37 °C in closed cap 15mL conical tubes to reach 10^8^ CFU/mL. Cells were pelleted, resuspended in 10mL of PBS, and bacterial cell density was verified by dilution and plating on TSA. Mice were then inoculated with 10^7^ CFU by retro-orbital injection of 100μL of bacterial suspension. At 48 hours post infection, antibiotic treatment was initiated at 400mg/kg of oxacillin and administered intraperitoneally every 12 hours over the course of three days. At the end of this treatment period, day five post infection, the kidney, heart, and livers were harvested and homogenized to determine bacterial burdens.

## ACKNOWLEDGMENTS

The presented research in this publication was 100% funded by the National Institute of Allergy and Infectious Diseases (NIAID) of the National Institutes of Health (NIH) under Award Numbers R01AI179695, R01AI173004 and R21AI159369. All program costs were covered by these grants and the content and views expressed in this manuscript are the sole responsibility of the authors, not necessarily reflective of the official views held by the NIH. The training of AZV was supported by NIAID/NIH Award Number F30AI169746. The Microscopy Services Laboratory, Department of Pathology and Laboratory Medicine, is supported in part by P30 CA016086 Cancer Center Core Support Grant to the UNC Lineberger Comprehensive Cancer Center. The Zeiss LSM900 microscope utilized in this project was funded with support from NIH Grant S10OD036215. The NTML Genetic Toolbox and transposon mutants were provided by the Network on Antimicrobial Resistance in *Staphylococcus aureus* (NARSA) for distribution by BEI Resources. We thank Kristen White and Jillann Madren for their help with TEM as well as Pablo Ariel and Ashelyn Sidders for their guidance in confocal microscopy. We thank Basilea Pharmaceutica for providing ceftobiprole and Robert Bourret for his helpful input and experimental guidance over the course of this project.

## REFERENCES

1. K. Bush, P. A. Bradford, β-Lactams and β-Lactamase Inhibitors: An Overview. Cold Spring Harb Perspect Med 6, (2016).

2. P. Piewngam, M. Otto, Staphylococcus aureus colonisation and strategies for decolonisation. The Lancet Microbe 5, e606–e618 (2024).

3. C. M. Pandey N, in Beta-Lactam Antibiotics. (StatPearls Publishing, Treasure Island (FL), Updated 2023 Jun 4).

4. T. L. Holland et al., Ceftobiprole for Treatment of Complicated Staphylococcus aureus Bacteremia. N Engl J Med 389, 1390–1401 (2023).

5. K. V. Bergersen et al., Early cytokine signatures and clinical phenotypes discriminate persistent from resolving MRSA bacteremia. BMC Infect Dis 25, 231 (2025).

6. M. Wang, G. Buist, J. M. van Dijl, Staphylococcus aureus cell wall maintenance - the multifaceted roles of peptidoglycan hydrolases in bacterial growth, fitness, and virulence. FEMS Microbiol Rev 46, (2022).

7. A. Vermassen et al., Cell Wall Hydrolases in Bacteria: Insight on the Diversity of Cell Wall Amidases, Glycosidases and Peptidases Toward Peptidoglycan. Frontiers in Microbiology 10, (2019).

8. T. Dörr, Understanding tolerance to cell wall-active antibiotics. Ann N Y Acad Sci 1496, 35–58 (2021).

9. K. Kitano, A. Tomasz, Triggering of autolytic cell wall degradation in Escherichia coli by beta-lactam antibiotics. Antimicrobial Agents and Chemotherapy 16, 838–848 (1979).

10. A. Tomasz, S. Waks, Mechanism of action of penicillin: triggering of the pneumococcal autolytic enzyme by inhibitors of cell wall synthesis. Proceedings of the National Academy of Sciences 72, 4162–4166 (1975).

11. J. P. Zackular, W. J. Chazin, E. P. Skaar, Nutritional Immunity: S100 Proteins at the Host-Pathogen Interface. J Biol Chem 290, 18991–18998 (2015).

12. S. M. Damo et al., Molecular basis for manganese sequestration by calprotectin and roles in the innate immune response to invading bacterial pathogens. Proc Natl Acad Sci U S A 110, 3841–3846 (2013).

13. E. M. Zygiel, E. M. Nolan, Exploring Iron Withholding by the Innate Immune Protein Human Calprotectin. Acc Chem Res 52, 2301–2308 (2019).

14. T. G. Nakashige, E. M. Zygiel, C. L. Drennan, E. M. Nolan, Nickel Sequestration by the Host-Defense Protein Human Calprotectin. J Am Chem Soc 139, 8828–8836 (2017).

15. A. N. Besold et al., Role of Calprotectin in Withholding Zinc and Copper from Candida albicans. Infect Immun 86, (2018).

16. B. D. Corbin et al., Metal chelation and inhibition of bacterial growth in tissue abscesses. Science 319, 962–965 (2008).

17. M. I. Hood et al., Identification of an Acinetobacter baumannii zinc acquisition system that facilitates resistance to calprotectin-mediated zinc sequestration. PLoS Pathog 8, e1003068 (2012).

18. T. E. Kehl-Fie et al., MntABC and MntH contribute to systemic Staphylococcus aureus infection by competing with calprotectin for nutrient manganese. Infect Immun 81, 3395–3405 (2013).

19. P.K. Párraga Solórzano, T. S. Bastille, J. N. Radin, T. E. Kehl-Fie, A Manganese-independent Aldolase Enables Staphylococcus aureus To Resist Host-imposed Metal Starvation. mBio 14, e0322322 (2023).

20. J. N. Radin et al., Metal-independent variants of phosphoglycerate mutase promote resistance to nutritional immunity and retention of glycolysis during infection. PLoS Pathog 15, e1007971 (2019).

21. Y. M. Garcia et al., A Superoxide Dismutase Capable of Functioning with Iron or Manganese Promotes the Resistance of Staphylococcus aureus to Calprotectin and Nutritional Immunity. PLoS Pathog 13, e1006125 (2017).

22. S. L. Price et al., Yersiniabactin contributes to overcoming zinc restriction during Yersinia pestis infection of mammalian and insect hosts. Proc Natl Acad Sci U S A 118, (2021).

23. L. R. Burcham et al., Identification of Zinc-Dependent Mechanisms Used by Group B Streptococcus To Overcome Calprotectin-Mediated Stress. mBio 11, (2020).

24. M. Nega, P. M. Tribelli, K. Hipp, M. Stahl, F. Götz, New insights in the coordinated amidase and glucosaminidase activity of the major autolysin (Atl) in Staphylococcus aureus. Commun Biol 3, 695 (2020).

25. J. N. Radin, J. Zhu, E. B. Brazel, C. A. McDevitt, T. E. Kehl-Fie, Synergy between Nutritional Immunity and Independent Host Defenses Contributes to the Importance of the MntABC Manganese Transporter during Staphylococcus aureus Infection. Infect Immun 87, (2019).

26. D. M. Bamberger, S. E. Boyd, Management of Staphylococcus aureus infections. Am Fam Physician 72, 2474–2481 (2005).

27. M. Wolkewitz, U. Frank, G. Philips, M. Schumacher, P. Davey, Mortality associated with in-hospital bacteraemia caused by Staphylococcus aureus: a multistate analysis with follow-up beyond hospital discharge. J Antimicrob Chemother 66, 381–386 (2011).

28. S. E. Rowe et al., Reactive oxygen species induce antibiotic tolerance during systemic Staphylococcus aureus infection. Nat Microbiol 5, 282–290 (2020).

29. J. E. Beam et al., Inflammasome-mediated glucose limitation induces antibiotic tolerance in Staphylococcus aureus. iScience 26, 107942 (2023).

30. S. Helaine, B. P. Conlon, K. M. Davis, D. G. Russell, Host stress drives tolerance and persistence: The bane of anti-microbial therapeutics. Cell Host Microbe 32, 852–862 (2024).

31. E. V. K. Ledger, S. Mesnage, A. M. Edwards, Human serum triggers antibiotic tolerance in Staphylococcus aureus. Nat Commun 13, 2041 (2022).

32. J. R. Raneses, A. L. Ellison, B. Liu, K. M. Davis, Subpopulations of Stressed Yersinia pseudotuberculosis Preferentially Survive Doxycycline Treatment within Host Tissues. mBio 11, 10.1128/mbio.00901-00920 (2020).

33. C. C. Murdoch, E. P. Skaar, Nutritional immunity: the battle for nutrient metals at the host–pathogen interface. Nature Reviews Microbiology 20, 657–670 (2022).

34. T. E. Kehl-Fie, E. P. Skaar, Nutritional immunity beyond iron: a role for manganese and zinc. Curr Opin Chem Biol 14, 218–224 (2010).

35. K. P. Grim et al., The Metallophore Staphylopine Enables Staphylococcus aureus To Compete with the Host for Zinc and Overcome Nutritional Immunity. mBio 8, (2017).

36. D. M. Vermilyea, A. W. Crocker, A. H. Gifford, D. A. Hogan, Calprotectin-Mediated Zinc Chelation Inhibits Pseudomonas aeruginosa Protease Activity in Cystic Fibrosis Sputum. J Bacteriol 203, e0010021 (2021).

37. A. J. Laarman et al., Staphylococcus aureus Metalloprotease Aureolysin Cleaves Complement C3 To Mediate Immune Evasion. The Journal of Immunology 186, 6445–6453 (2011).

38. M. Firczuk, M. Bochtler, Folds and activities of peptidoglycan amidases. FEMS Microbiology Reviews 31, 676–691 (2007).

39. M. Modi, M. Thambiraja, A. Cherukat, R. M. Yennamalli, R. Priyadarshini, Structure predictions and functional insights into Amidase_3 domain containing N-acetylmuramyl-L-alanine amidases from Deinococcus indicus DR1. BMC Microbiol 24, 101 (2024).

40. A. Usenik et al., The CWB2 Cell Wall-Anchoring Module Is Revealed by the Crystal Structures of the Clostridium difficile Cell Wall Proteins Cwp8 and Cwp6. Structure 25, 514–521 (2017).

41. A. Razew, J. N. Schwarz, P. Mitkowski, I. Sabala, M. Kaus-Drobek, One fold, many functions-M23 family of peptidoglycan hydrolases. Front Microbiol 13, 1036964 (2022).

42. C. A. Lopez, E. P. Skaar, The Impact of Dietary Transition Metals on Host-Bacterial Interactions. Cell Host Microbe 23, 737–748 (2018).

43. L. J. Juttukonda et al., Dietary Manganese Promotes Staphylococcal Infection of the Heart. Cell Host Microbe 22, 531-542.e538 (2017).

44. J. L. Bose, P. D. Fey, K. W. Bayles, Genetic tools to enhance the study of gene function and regulation in Staphylococcus aureus. Appl Environ Microbiol 79, 2218–2224 (2013).

45. T. E. Kehl-Fie et al., Nutrient metal sequestration by calprotectin inhibits bacterial superoxide defense, enhancing neutrophil killing of Staphylococcus aureus. Cell Host Microbe 10, 158–164 (2011).

46. A. E. Sidders et al., Antibiotic-induced accumulation of lipid II synergizes with antimicrobial fatty acids to eradicate bacterial populations. eLife 12, e80246 (2023).

