## Supplemental Information for "The innate immune protein calprotectin incapacitates the bactericidal activity of β-lactam antibiotics"

**SUPPLEMENTARY INFORMATION**

**Supplementary Table 1. Strain list**

| Strain | Genotype/strain description | Citation/source |
|---|---|---|
| *S. aureus* Newman | Wild-type | (*1*) |
| *S. aureus* JE2 | Wild-type | (*2*) |
| *S. aureus* Δatl/ΔlytM | Atl::Tn (erm) lytM::Tn (kan) Newman | Current study |
| *S. aureus SA03803* | Clinical isolate (MSSA) | Current study/SABG* |
| *S. aureus SA03833* | Clinical isolate (MRSA) | Current study/SABG* |
| *S. aureus SA03850* | Clinical isolate (MSSA) | Current study/SABG* |

*Duke University Hospital, *S. aureus* Bacteremia Group (SABG) biorepository

**Supplementary Table 2. The addition of calprotectin does not change the antibiotic minimum inhibitory concentration (MIC) of select β-lactams.** *S. aureus* Newman was incubated in 2-fold increasing concentrations of cefazolin, oxacillin, and nafcillin in the presence or absence of 240 μg/mL CP. The MIC was determined by visual identification of the concentration showing absence of bacterial growth after 24-hour incubation at 37 °C.

| Strain | Antibiotic | MIC | MIC +CP240 |
|---|---|---|---|
| *S. aureus* Newman | cefazolin | 0.25μg/mL | 0.25-0.5μg/mL |
| *S. aureus* Newman | oxacillin | 0.125μg/mL | 0.125-0.25μg/mL |
| *S. aureus* Newman | nafcillin | 0.25-0.5μg/mL | 0.25μg/mL |

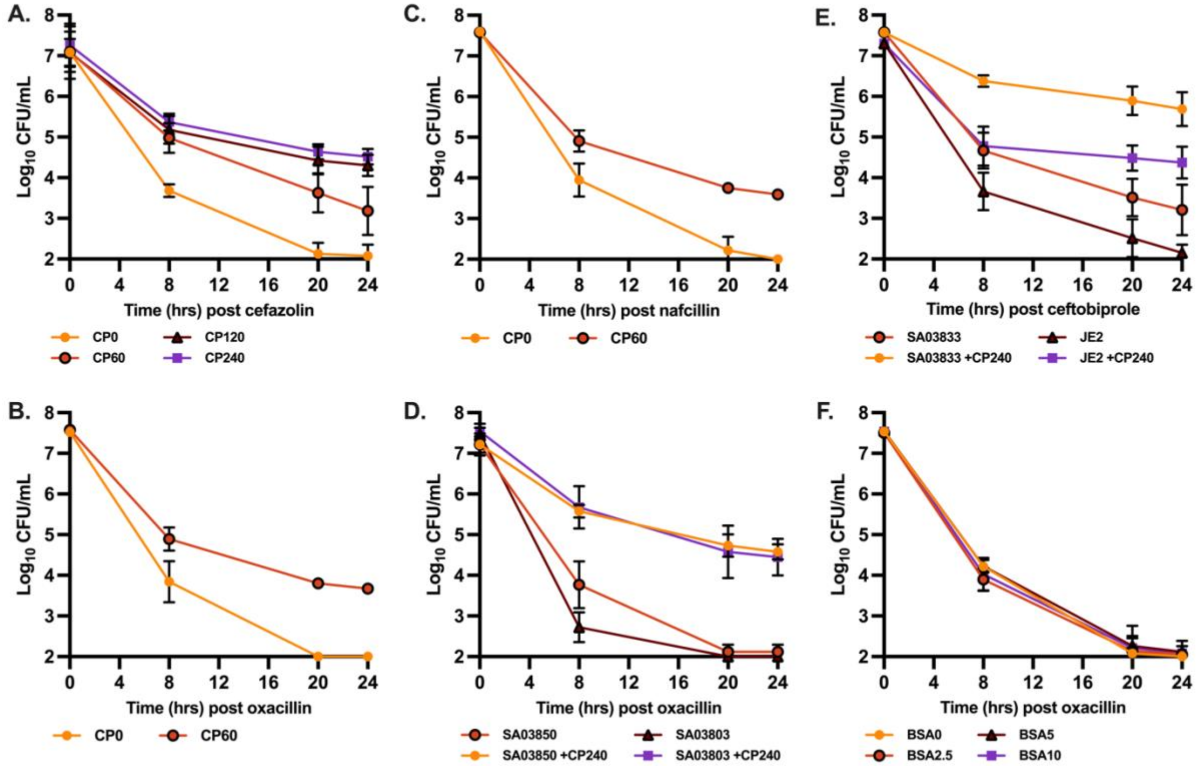


**Supplementary Figure S1. The presence of calprotectin negatively impacts the efficacy of cell-wall acting antibiotics against multiple MSSA and MRSA clinical and laboratory strains.** *S. aureus* Newman was grown to mid-exponential phase in TSB based media. Following determination of the starting bacterial burden (0hr), cultures were treated with (A) cefazolin at 50 μg/mL and 0-240 μg/mL CP; (B) oxacillin at 50 μg/mL and 0-60 μg/mL CP; or (C) nafcillin at 50 μg/mL and 0-60 μg/mL CP. *S. aureus* clinical isolates were grown to mid $10^7$ CFU/mL in in TSB based media. (D) MSSA clinical isolates (SA03850 and SA03803) were challenged with oxacillin at 50 μg/mL and 240 μg/mL CP. (E) *S. aureus* MRSA clinical isolate (SA03833) and laboratory isolate JE2 were challenged with ceftobiprole at 30 μg/mL and 240 μg/mL CP. *S. aureus* Newman was grown to mid $10^7$ CFU/mL in in TSB based media and cultures were then treated with (F) oxacillin at 50 μg/mL and 0-10 μM of Bovine Serum Albumin (BSA). For all experiments, bacterial burden at 8 hr, 20 hr, and 24 hr post treatment was determined by dilution plating. Data graphed shows the mean and standard deviation for experiments completed on three separate days each using two independent cultures (N=6).

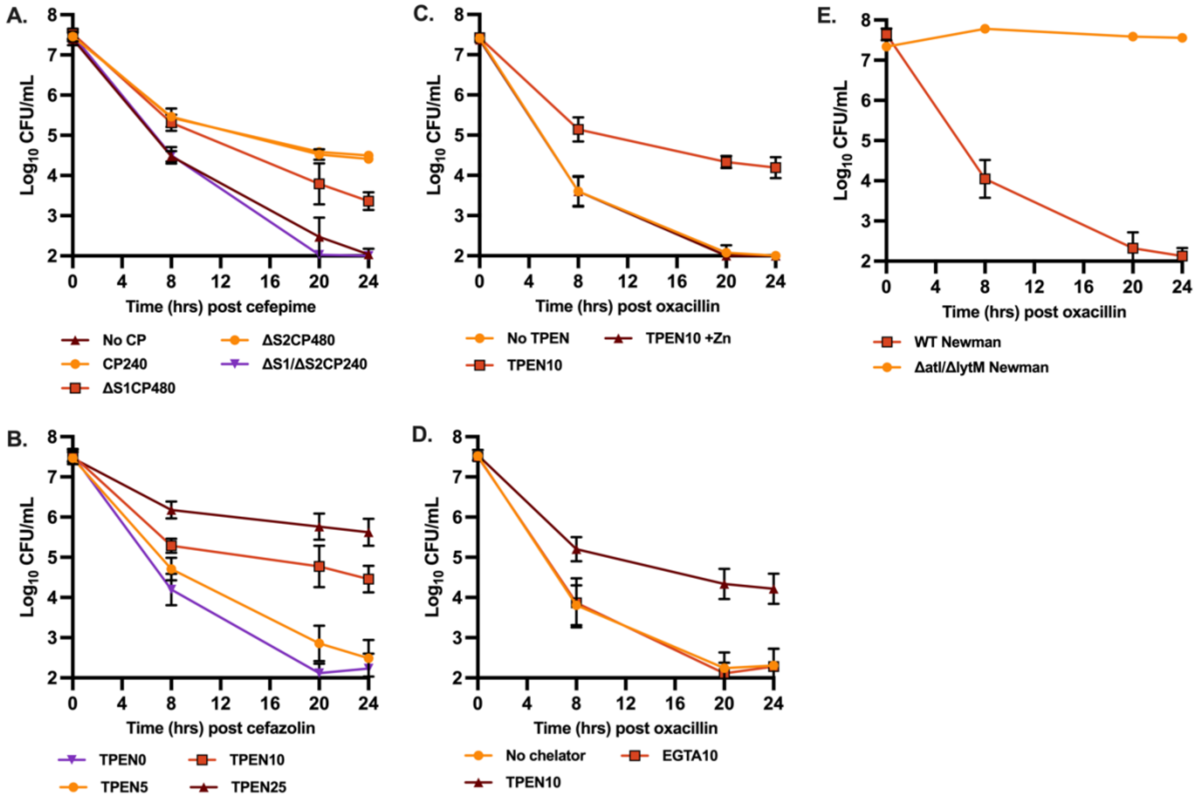


**Supplementary Figure S2. Chelation of zinc or the loss of autolytic activity, induces antibiotic tolerance.** *S. aureus* Newman was grown to mid-exponential phase in TSB based media. Following determination of the starting bacterial burden (0 hr), cultures were treated with (A) cefepime at 30 μg/mL and wildtype CP (240 μg/mL), ΔS1/ΔS2 (240 μg/mL), ΔS1 (480 μg/mL), or ΔS2 (480 μg/mL); (B) cefazolin at 50 μg/mL with TPEN (5–25 μM); (C) oxacillin at 50 μg/mL and 10 μM TPEN with and without the addition of 50 μM $ZnSO_4$; or (D) oxacillin at 50 μg/mL and 10 μM TPEN or 10 μM EGTA. (E) *S. aureus* Newman wildtype or Δ*atl/lytM* was treated with oxacillin at 50 μg/mL. For all experiments, bacterial burden at 8 hr, 20 hr, and 24 hr post treatment was determined by plating on TSA to enumerate survivors. Data graphed shows the mean and standard deviation for experiments completed on three separate days each using two independent cultures (N=6).

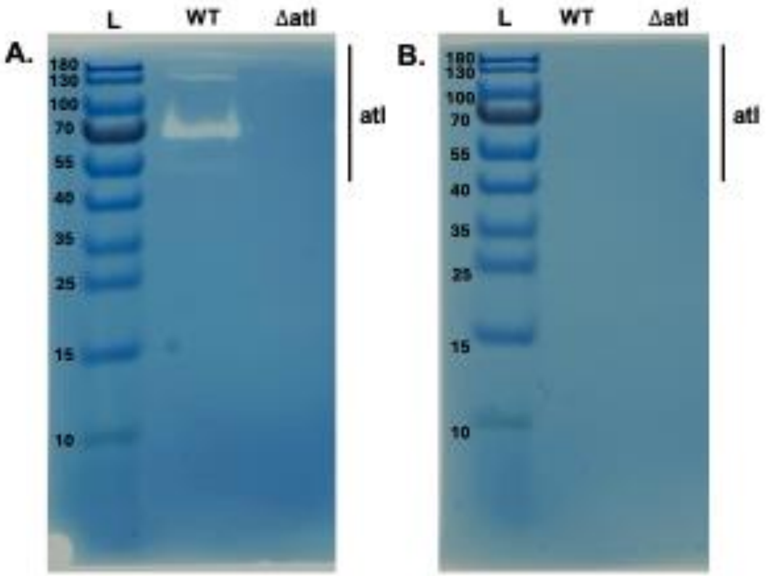


**Supplementary Figure S3. TPEN decreases Atl signal in zymogram.** Cell wall-associated proteins were isolated from 5 mL of overnight JE2 cultures, both wildtype and Δ*atl* JE2. Supernatant autolysins were then separated by SDS-PAGE alongside a protein ladder for size determination. 10% resolving gels were cast containing 2 mg/mL of heat-killed *S. aureus* RN4220, in the (A) absence or (B) presence of 240 μM TPEN. A representative image from one independent experiment is presented and two independent sets of gels were analyzed showing equivalent results.

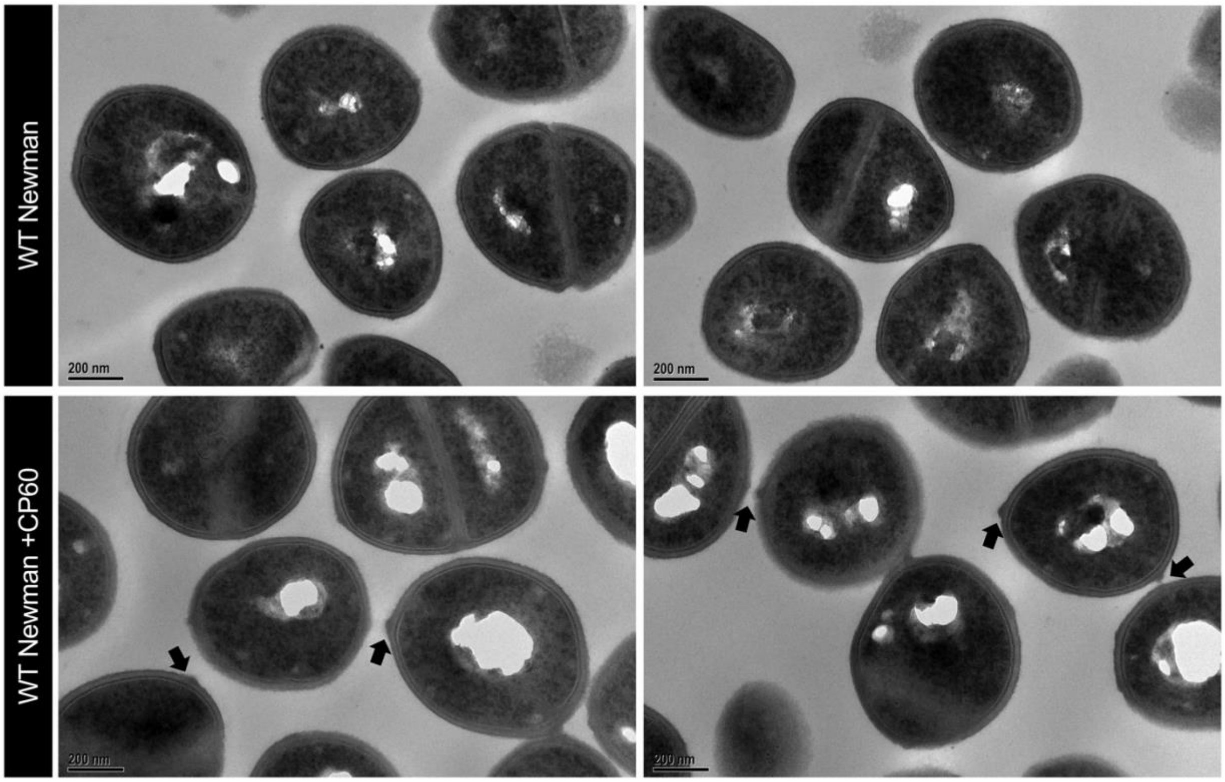


**Supplementary Figure S4. Changes to the structure of the cell wall in the presence of calprotectin.** Cell wall structure post 2-hour incubation with 60 μg/mL CP was visualized using TEM at an imaging magnification of 100,000X. Black arrows within the CP-treated condition highlight regions of peptidoglycan accumulation.

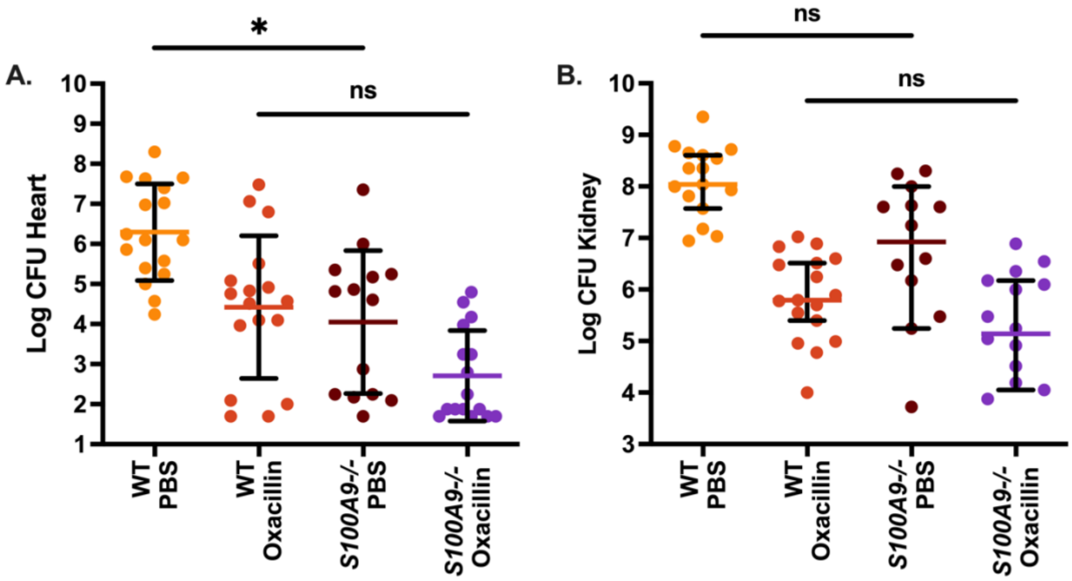


**Supplementary Figure S5. Bacterial burden of *S. aureus* within heart and kidney tissue comparing untreated and oxacillin treated wild-type and S100A9$^{-/-}$ mice.** Wild-type and S100A9$^{-/-}$ C57BL/6 mice were infected retro-orbitally with $10^7$ CFU Newman and treated with oxacillin. After three days of treatment, the bacterial burden in the heart (A) and kidneys (B) was determined. Each data point represents one individual mouse and graphed is the combination of two independent experiments. Data graphed shows the median with 95% CI. Statistical significance ($p \leq 0.05$) was determined by Kruskal-Wallis with Dunn's multiple comparisons.

### METHODS

**Minimum inhibitory concentration determination:** The minimum inhibitory concentration (MIC) in the presence of CP was determined using a broth antibiotic microdilution assay for cefazolin, oxacillin, and nafcillin. Antibiotics were diluted 2-fold from 4 μg/mL down to 0.03 μg/mL in a 96-well plate. Overnight cultures of *S. aureus* Newman were then diluted 1:1000 to reach around $5 \times 10^6$ CFU/mL and combined with calprotectin to a final concentration of 240 μg/mL. The MIC was determined by visual observation of growth inhibition following a 24-hour incubation at 37 °C.

**Zymography analysis:** Overnight cultures of *S. aureus* JE2 grown in 5 mL of TSB were pelleted and washed twice in PBS. After the final wash, cells were resuspended in 1 mL of 4% SDS. After 45 minutes gentle rotation at room temperature, cells were spun down at 8000 rpm for 5 min, and the supernatant containing autolysins was collected. 100 uL aliquots were then treated with 1.2 mM TPEN or left untreated and incubated for an additional 30 min. Zymogram gels were constructed similar to what has been previously described (*3*). For our experiments, a 10% resolving gel was cast containing 2 mg/mL of heat-killed *S. aureus* RN4220, with and without the addition of 240 μM TPEN. Supernatant autolysins were then separated by SDS-PAGE alongside a protein ladder for size determination. Gels were then incubated overnight in renaturation buffer (50 mM tris HCl, 0.1% Triton x-100, 10 mM $CaCl_2$, 10 mM $MgCl_2$) and stained with 0.1% methylene blue dissolved in 0.01% KOH. Gels were imaged using a light pad to visualize regions of clearance.

**Transmission Electron Microscopy:** Bacterial cell pellets were fixed in 2% paraformaldehyde/2.5% glutaraldehyde in 0.1 M sodium phosphate buffer, pH 7.4 for 1 hour at room temperature. Cell pellets were washed three times for ten minutes each at room temperature in 0.1 M sodium phosphate buffer, pH 7.4. A solution of 1% osmium tetroxide buffered in 0.15 M sodium phosphate was added to each sample and allowed to sit at room temperature for one hour. Cells were then washed three times with deionized water and dehydrated with an increasing ethanol gradient with en bloc uranyl acetate staining performed for 1 hour with the 50% ethanol (30%, 50% with 2% uranyl acetate 1 hour, 75%, 90%, 100%, 100%, 100%, 10 minutes each at room temperature) followed by two rounds of propylene oxide for 30 minutes each. Cell pellets were then infiltrated with 1:1 propylene oxide:Spurr’s resin overnight, exchanged with 100% Spurr’s resin for six hours, exchanged with 100% Spurr's once more and cured at 60 °C overnight. Ultrathin sections (80 nm) were cut using a diamond knife on a Leica UCT7 ultramicrotome and applied to 200 mesh copper grids. Grids were stained with

4% aqueous uranyl acetate for 12 minutes followed by Reynold's lead citrate for 8 minutes (*4*). Samples were viewed using a JEOL JEM-1230 transmission electron microscope operating at 80 kV (JEOL USA, Inc., Peabody, MA) and images were acquired with a Gatan Orius SC1000 CCD Digital Camera and Gatan Microscopy Suite 3.0 software (Gatan, Inc).

### REFERENCES


1. J. N. Radin, J. Zhu, E. B. Brazel, C. A. McDevitt, T. E. Kehl-Fie, Synergy between Nutritional Immunity and Independent Host Defenses Contributes to the Importance of the MntABC Manganese Transporter during Staphylococcus aureus Infection. *Infect Immun* **87**, (2019).
2. P. D. Fey, J. L. Endres, V. K. Yajjala, T. J. Widhelm, R. J. Boissy, J. L. Bose, K. W. Bayles, A genetic resource for rapid and comprehensive phenotype screening of nonessential Staphylococcus aureus genes. *mBio* **4**, e00537-00512 (2013).
3. I. Thalsø-Madsen, F. R. Torrubia, L. Xu, A. Petersen, C. Jensen, D. Frees, The Sle1 Cell Wall Amidase Is Essential for β-Lactam Resistance in Community-Acquired Methicillin-Resistant Staphylococcus aureus USA300. *Antimicrob Agents Chemother* **64**, (2019).
4. E. S. Reynolds, The use of lead citrate at high pH as an electron-opaque stain in electron microscopy. *J Cell Biol* **17**, 208-212 (1963).
